# Semi-Automated Seal Detection on the Western Antarctic Peninsula: An Unsupervised Machine Learning Approach for Detecting Ice Seals in Aerial Survey Data

**DOI:** 10.1101/2025.08.22.671691

**Authors:** C. McGinnity, C.C.G. Bamford, N. Fenney, A. Fleming, J. Forcada, M.S. Tift, L.A. Hückstädt, D.P. Costa, P.T. Fretwell

## Abstract

Over the past 25 years, the Western Antarctic Peninsula (WAP) has experienced dramatic shifts in sea ice extent. This change has coincided with rapid alterations in ice-dependent ecosystems, including those supporting crabeater seals - the most abundant Antarctic seal and one of the largest mammalian consumers of krill. Despite their ecological importance, population estimates for ice seals remain scarce due to the difficulty of surveying large-scale, remote, ice-covered habitats. In 2023, during an abnormally low sea ice year, we conducted aerial surveys over Crystal Sound and Marguerite Bay during the end of the breeding season, flying over 1,000 km of transects. Seals were extremely sparse in the resulting imagery - occupying less than 1% of the surveyed area. This posed a significant challenge for both manual annotation and automated detection. Here we present a semi-automated, rules-based image analysis pipeline to substantially reduce human annotation time. Our method leverages hierarchical clustering with just two tunable parameters, avoiding the computational burden and opacity of deep learning models. Using this method, we identified 758 seals within a ∼350 km^2^ survey subset, achieving a test recall of 82%. In the absence of concurrent tagging data to estimate haul-out corrections, we refrain from extrapolating to a population estimate. However, the low observed densities highlight the urgent need for continued monitoring. Our improved data processing pipeline is a key step in facilitating the large-scale analysis required to inform conservation strategies for this key species.

## I. Introduction

Since the peak in Antarctic sea ice extent in 2014, subsequent years have been marked by dramatic, non-linear declines [1, 2, 3] that deviate from IPCC projections [4], occurring alongside atmospheric warming far exceeding the global average [5, 6, 7, 8, 9], causing profound impacts on the physical environment of the Antarctic Peninsula.

In terms of sea ice coverage and seasonality, sea surface temperatures have risen [10, 11, 12]; seasonal sea ice thickness has decreased, and its persistence has shortened by three months [13, 8]. Glaciers have thinned [6], freshening coastal waters [14, 15], which in turn has triggered biological responses. Notably, there has been a shift toward gelatinous zooplankton assemblages [16], along with a southward redistribution of Antarctic krill (*Euphausia superba*; hereafter, krill) biomass [17, 18, 19]. There has also been a significant reduction in suitable habitat—particularly pack ice—for ice-dependent penguin and seal species [20, 21, 22].

Among the ice-dependent seal species on the Western Antarctic Peninsula (WAP), crabeater seals (*Lobodon carcinophaga*) are by far the most abundant, with an estimated population exceeding 1.8 million in the WAP [20], the highest known densities of this species anywhere on the continent. Despite their name, crabeater seals rely on krill for >88% of their diet [23], making them a key indicator species [24] as one of Antarctica’s most specialised foragers and one of the most important air breathing consumers of krill [25, 26, 27, 23]. However, data collection on crabeater seals is challenging due to their year-round residence within the pack ice [28, 20], a notoriously difficult habitat to sample without disturbance, and the fact that they do not aggregate at breeding colonies nor haul out at predictable locations. Crabeater seals are also sighted at sea, and in large numbers, adding to the complexity of data acquisition. As such, our understanding of their population is coarse and out of date, an issue amplified by the fact that the ice conditions on the WAP have changed rapidly in recent years.

Population metrics are essential to inform and validate theoretical models, which in turn guide management and conservation strategies [29]. Remote sensing overcomes many of the traditional difficulties of surveying pack ice species by ship [30], but presents new challenges, namely the processing of the drastically increased volume of data. Satellites and aircraft (both piloted and remotely operated) are the main platforms currently used for animal detection [31]. However, these platforms, which are often photo-based, produce data volumes that are impractical to process manually [32, 33]. To overcome processing demands, researchers have implemented various methods including citizen science campaigns to perform a “search area reduction” of >260,000km^2^ of fast ice [34] and developing Convolutional Neural Networks (CNN) to detect pack ice seals [35], wildebeest (*Connochaetes taurinus/gnou*) [36], and whales [37]. However, Deep Learning approaches such as CNNs inherently lack transparency, interpretability and explainability due to the model structure [38] and may fail to generalize [36] outside their original, often narrow, training environment. This may in part have led to low uptake of such methods, particularly with the aforementioned species and within the ecological field.

Deep learning is a powerful tool but requires large volumes of labelled training data that are both specific to and representative of the survey environment [39]. Published models are often highly case-specific and typically require retraining for each new application [36]. Another major challenge is the scarcity of publicly available training datasets for wildlife detection [31]; those that do exist are often narrowly focused on a particular species, geographic region, or data-type [40]. Consequently, deploying deep learning in new conditions demands extensive and tedious manual data labelling, a process which is widely acknowledged not to produce perfect ground truth on which to train and evaluate a model, with errors in the training labels significantly reducing the accuracy of neural net classifiers [41].

An especially challenging attribute of aerial surveys for pack-ice seals is the sparsity of targets (*i*.*e*., ice seals) within the vast background. The categorisation of binary classification tasks for which a single class represents <15% of the distribution is regarded as highly unbalanced [42]. However, for seals in pack ice environments, individuals are found to occupy <1% [43, 20] of the background [43], a trend similarly reflected in our data (Figure 1).

**Figure 1.**
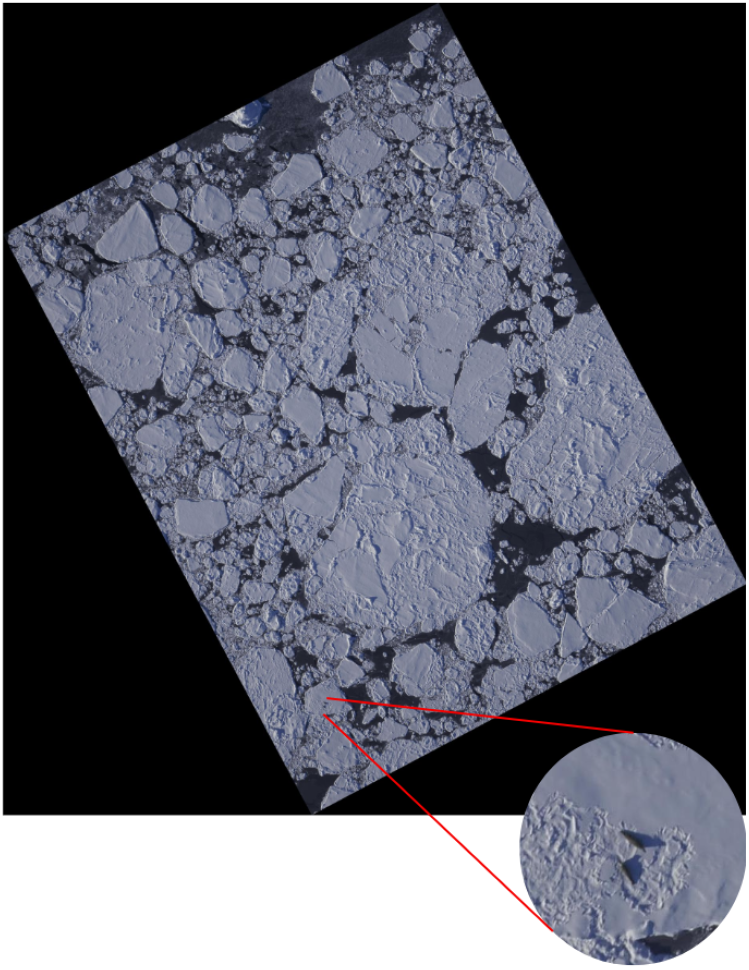
Example of a single aerial image captured during the 2023 survey of the Western Antarctic Peninsula (WAP). Seals occupy less than 1% of the surveyed area, posing a significant challenge for classification algorithms as imbalanced data sets often skew predictive performance toward the majority class, increasing the risk of seals going undetected.

Our classification problem is therefore a case of extreme imbalance, which presents challenges for algorithms and human annotators alike. Imbalance is known to bias the predictive performance of many classification algorithms toward the majority class [44], which in our case would be negative. Manual annotation suffers from similar performance problems, for instance, Rožanec et al. measured a 3 fold increase in precision of human annotators when labelling a balanced dataset over an imbalanced set, linking this to improved participant attention [45]. Additionally, the quality of human labelling is known to degrade over time with fatigue [46], which presents a further concern for the processing of vast quantities of data produced in wildlife surveying.

In the face of the challenges posed by data sparsity, class imbalance, and the limitations of both deep learning and human annotation, we adopt a parsimonious approach to analyse data from a set of aerial surveys flown over Crystal Sound and Marguerite Bay in 2023 (Figure 2). Rules based algorithms have been successfully deployed to significantly reduce researcher workload. For example, the count time for humpback whales was reduced from from 24 hours to ∼30 mins by filtering the satellite data using just a shape criteria [47]. Building on this rationale, we present a semi-automated, rules-based image analysis pipeline that leverages unsupervised machine learning, using hierarchical clustering to streamline data processing with minimal parameter tuning to significantly reduce the manual annotation burden, making the processing of regular and large-scale wildlife surveys more feasible.

**Figure 2.**
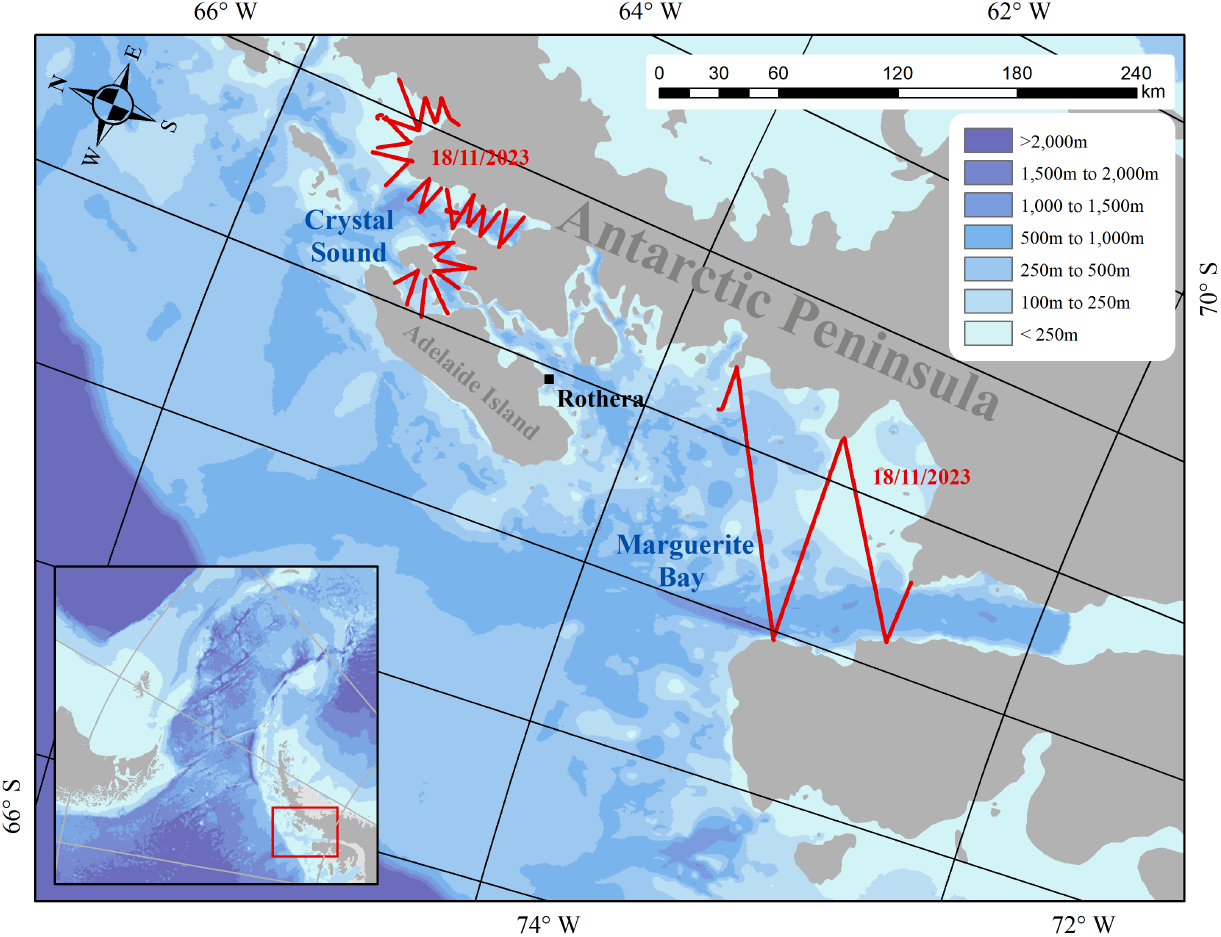
Surveys flown on the 18^th^ November 2023 over Crystal Sound (560.2 km) and Marguerite Bay (481.4 km). Flights were conducted using a twin-otter aircraft equipped with a Phase One iXU 150 medium format aerial camera (3-band, RGB). These regions were selected based on the typical prevalence of pack-ice and the known association of the target species with this habitat.

In this study, we apply such methods to introduce and evaluate a semi-automated image processing pipeline. We then review previous seal survey efforts on the WAP, highlighting methodological inconsistencies that have hindered long-term comparisons. Finally, we argue for standardised survey design and concurrent tagging efforts as essential components of a sustainable monitoring strategy. Together, these steps offer a practical framework for tracking population trends in a dynamic polar ecosystem.

## II. Materials AND Methods

### i. Data Collection & Aerial Photography

In 2023, dedicated surveys were flown with a De Havilland Canada DHC-6 Series 300 Twin Otter aircraft operated by the British Antarctic Survey out of Rothera Research Station (67.6°S 68.1°W, Figure 2). These flights took place in mid-November at the end of the breeding season and targeted pack ice habitat typically associated with crabeater seals, with the chosen embayments anecdotally known for high densities of this species [20]. However, 2023 was a year of record sea ice lows with a maximum extent 10% below the average and a shortening of the sea ice season by 3-4 months in most regions [48]. It is unknown how this sea ice volatility may have impacted ice seals in the surveyed region as reference baselines are limited.

Data were collected using a Phase One iXU 150 medium format aerial camera (3-band, RGB), with a sensor resolution of 50 megapixels and a 55 mm Schneider lens. The surveys were undertaken at a target altitude of 500 m, aiming to achieve a ground sampling distance (GSD) of 4.5 cm and a swath width of 400 m. A total of 1041.6 km was flown over Crystal Sound and Marguerite Bay on the 18^th^ November 2023 (Figure 2). The Phase One iXU 150 was configured to capture an image every second, producing a forward overlap of around 80%. The camera system lacked an integrated IMU (inertial measurement unit), having only a GNSS (Global Navigation Satellite System) receiver to record the position of the camera in space. Without an IMU, collecting precise orientation information of the system during image capture was not possible. Instead, the average heading for each flight line was calculated and used to correct the orientation of the imagery. Using Socet GXP, each image was then orthorectified against a flat surface with an elevation of 0 m and exported as a georeferenced.TIF, using a geographic coordinate system (EPSG:4326 - WGS 84). A subset of the orthorectified image tiles spanning ∼350 km^2^ of ice were then passed to a semi-automated pipeline, described below, aimed to improve the efficiency of this process.

### ii. Seal Detection

Seals in each image tile were identified by applying an adaptation of a pipeline developed to process aerial magnetic surveys [49]. This process leverages real world knowledge and unsu-pervised machine learning to drastically reduce human review time while also improving recall by significantly mitigating the imbalance problem. We apply this new method to process >5,500 images from the surveys flown on 18^th^ November 2023.

Of the three main species of ice seals on the WAP each of them typically grow to lengths in excess of 2m, whether that be crabeater (2.05-2.4m; [50],[51]), Weddell (*Leptonychotes weddellii*, 2.5-3.3m; [52]) or leopard (*Hydrurga leptonyx*, 3.3-3.8; [53]). Their dark pigmentation contrasts with the largely homogeneous Antarctic environment making these species easily identifiable in aerial imagery. Initial experiments determined that the contrast between the seals and ice was most significant in the blue channel of the RGB imagery, with their low saturation in the Hue Saturation Value (HSV) decomposition aiding their differentiation from shadows cast by geological features. This is intuitive given the colour profile of the Antarctic environment.

Our method capitalises on these key priors, producing a model with just two tuneable parameters: (i) a darkness threshold on the blue channel of the RGB image and (ii) a threshold on the saturation channel of the HSV decomposition. This approach drastically reduces the necessary amount of training data, mitigates the risk of overfitting, and has improved interpretability compared to a CNN model.

Our model consists of five main stages (Figure 3): (a) data preprocessing, (b) contouring, (c) HSV conversion, (d) filtering and (d) clustering. The original image is blurred using a 2×2 kernel to reduce noise. In the contouring stage, the model removes areas that satisfy the darkness threshold but are obviously too small or large to be a crabeater seal. Areas smaller than 1m in both dimensions or larger than 5m in either are discarded. This eliminates large bodies of water and large portions of brash ice. To further eliminate irrelevant dark areas that meet the shape criteria but are not seals, we filter out regions of high saturation, typical of shadows cast on the ice by terrain.

**Figure 3.**
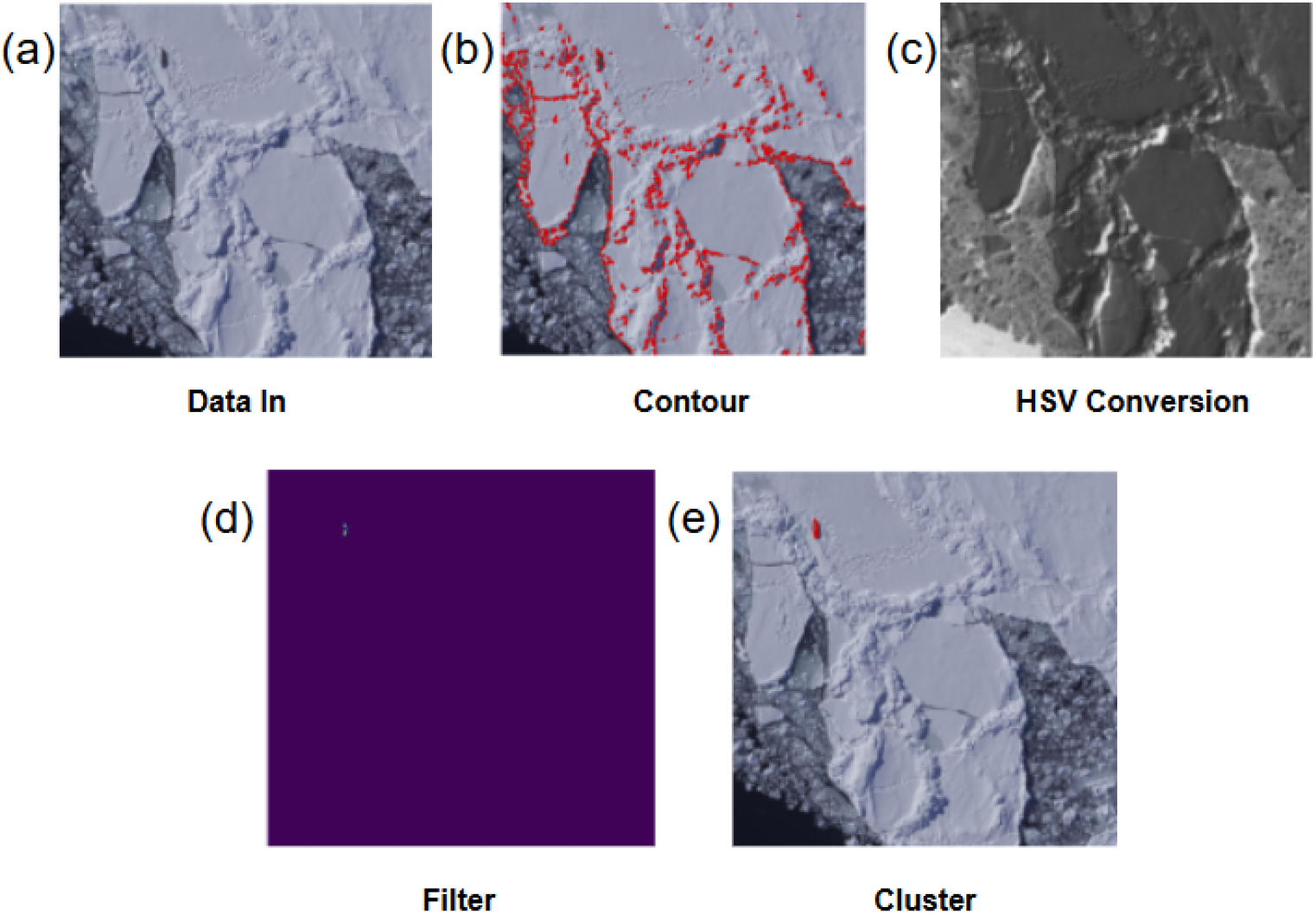
The pipeline exploits visual priors related to seal size, shape, and colour to streamline detection. The five main stages of our model are: (a) The original data is blurred with a 2×2 kernel to reduce noise. (b) Contouring eliminates large bodies of dark water and other irrelevant dark areas that do not match the seal shape criteria. (c) Seals are distinguished by their low saturation compared to other dark regions, such as shadows. (d) The applied filter thresholds reflect the selected parameters. (e) Remaining pixels are grouped into potential seals using hierarchical clustering.

Hierarchical clustering is then performed on the remaining pixels that satisfy the darkness threshold, using the single linkage method with distance metric. Clusters made up of less than 20 pixels, (<0.05m^2^) are immediately discarded as noise and the remaining are processed into a series of clipped images, centred on potential seals, ready for human review.

Figure 4 shows examples of the output of the automatic pipeline. Alongside the close up image, the reviewer is presented with a wider view to employ context clues, such as presence of tracks or ice holes, when performing the final classification. This also acts to catch and remove duplicated seals present due to the forward overlap in the aerial imagery. Even after tuning our parameters, we encounter a set of images, typically of complex fields of brash ice, that our algorithm fails to handle satisfactorily, flagging an excess number of potential seals. We subject these outlier images to traditional manual review, removing the top 5% of images ranked by total number of proposed seals.

**Figure 4.**
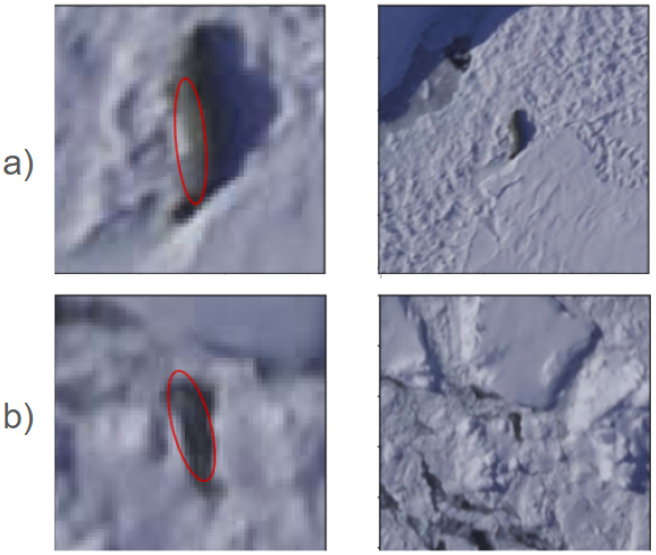
Example outputs from the semi-automated detection pipeline, showing cropped candidate seals alongside wider context imagery. This dual-view format allows the human reviewer to distinguish seals from similar features (e.g., shadows or terrain artifacts). Image (b) highlights a false positive, illustrating how terrain elements can resemble seals and the importance of contextual review in improving classification accuracy.

### iii. Model Tuning

In a effort to estimate a ground truth to enable evaluation of model performance, 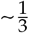 of the images were first manually reviewed by a human expert. The unaided reviewer identified 125 seals across ∼1600 images, with a review time in excess of 25 hours. Additionally, a second expert performed a subset review of ∼800 of these images, aided by a coarse filter, and detected 63 additional seals missed in the initial review.

We took clippings around these 188 seals and supplemented them with representative background images, creating a thinned dataset which was then divided into a training and test set with a 70:30 split. We performed a parameter grid search to tune our two parameters, and we computed the recall (Equation 1; proportion of actual positive cases correctly identified by the model) and precision (Equation 2; the proportion of predicted positive cases that are actually correct) of the automatic portion of our pipeline for all combinations of the saturation and darkness thresholds.

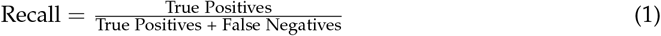

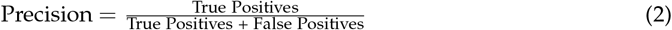

To select the final thresholds, we quantify the trade-off between recall and precision of the candidate values by computing the *F*_*β*_ score [54] for a range of *β* (Equation 3).

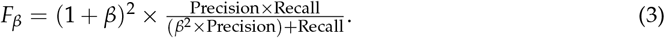

The thresholds that maximize this score are optimal for a given *β*. Here, *β* controls the relative weight of recall to precision, with higher values emphasizing recall. Importantly, *β* is not treated as a tuning parameter in the optimization process. Instead, we compute *F*_*β*_ scores across a range of *β* values to explore the trade-offs between recall and precision, and to illustrate the resource implications of different operating points. While *β* = 4 yields higher recall, it also results in a significantly higher human review burden. *β* is a design choice dependent on both the desired application of the model and the available resources.

### iv. Species identification and density estimation

Species identification of the processed image was conducted by an experienced observer who used differences in body shape, length, colouration, presence of aggregations and associated habitat type/feature to differentiate species (Table 1). Only ice obligate seals were classified to species level, with any ice-tolerant species, namely Antarctic fur (*Arctocephalus gazella*) and southern elephant seals *(Mirounga leonina*), were pooled with those where the clarity of the imagery prevented identification. Ice seal counts were used to estimate density (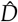) by dividing the number of observed seals by the ice area surveyed according to Equation 4. The total number of pixels was summed over the ∼5,500 processed images, excluding pixels corresponding to bodies of water or other non-ice areas. No adjustments were made for forward overlap between images in the density estimation. Instead, the total number of detected seals, divided by the total image area - both without accounting for forward overlap - is assumed to approximate the non-overlapping density estimate.

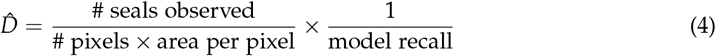

**Table 1.** Morphological and contextual characteristics used to distinguish between Antarctic ice seal species (crabeater, weddell, and leopard seals) in aerial imagery. Traits include body size and shape, colouration, group behaviour, and proximity to habitat features. These distinctions support manual species classification following initial automated detection.

| Species | Crabeater | Weddell | Leopard |
| --- | --- | --- | --- |
| Length | 2.05-2.4m | 2.5 - 3.3m | up to 3.8m |
| Body shape | Slender; pointed snout;<br>obvious scarring | Bulky fusiform shape | Elongated; bulky |
| Colouration | Medium brown to silver,<br>fading through the year | Dark, almost black with<br>grey streaks; spotted belly | Grey/silver |
| Aggregations | Solo to large aggregation possible | Small groups | Rare |
| Habitat cues | - | Proximity to breathing<br>holes | - |
| Sources | [50] | [52] | [53] |

As with all surveys of a species that are influenced by availability bias [55], the number of seals observed on the ice represents only a subset of the total number present. Accurate population estimates therefore require correction for the proportion of animals that are not hauled out at the time of the survey. This is typically achieved using bio-logging data to parameterise haul-out probabilities [20]. In the absence of tagging data for the season (spring), we do not attempt such extrapolation here.

## III. Results

### i. Model development

The recall and precision response surfaces for varying thresholds are shown in Figure 5. Our final model achieved a recall of 82%, surpassing the ∼50% recall from unaided manual review of the 2023 data, while also reducing the survey area requiring manual review by ∼94%.

**Figure 5.**
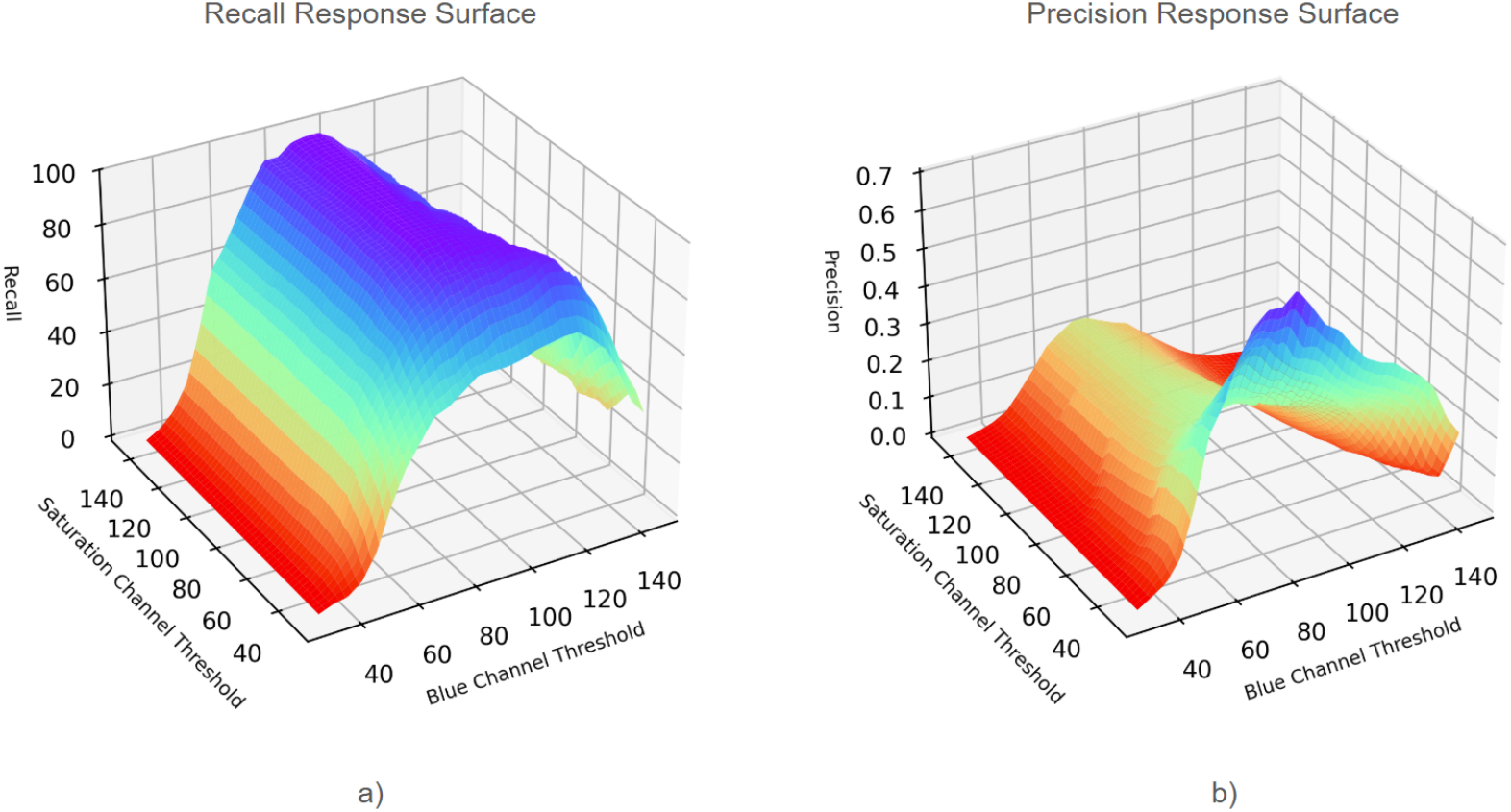
Response surfaces illustrate how model recall (a) and precision (b) vary as a function of threshold combinations applied to the blue channel and saturation channel of the RGB aerial imagery. The irregular shape of these surfaces indicate a non linear relationship between the parameters. Due to this non-linearity, strictly increasing a threshold is not guaranteed to capture more seals, therefore, thresholds were varied systematically across two dimensions, with a coarser grid in regions of low performance and a denser grid in regions approaching optimality. N.B. Colour-scale only serves as a visual aid.

The non linear relationship between the threshold parameters (Figure 5) indicates that relaxing a threshold does not necessarily result in the detection of more seals. The scatter plot of precision and recall for all threshold combinations (Figure 6) provides insight into the trade-off between these metrics. The final threshold choice will fall along the red frontier, with combinations beneath the curve being strictly suboptimal. The frontier’s gradient captures how quickly the recall declines while seeking greater precision. This provides a visual understanding of the effect of changing the value of *β*. The F score response surface reveals the optimal threshold choice for a given *β* as the surface global maximum. Figure 6 shows the response surface when *β* = 4.

**Figure 6.**
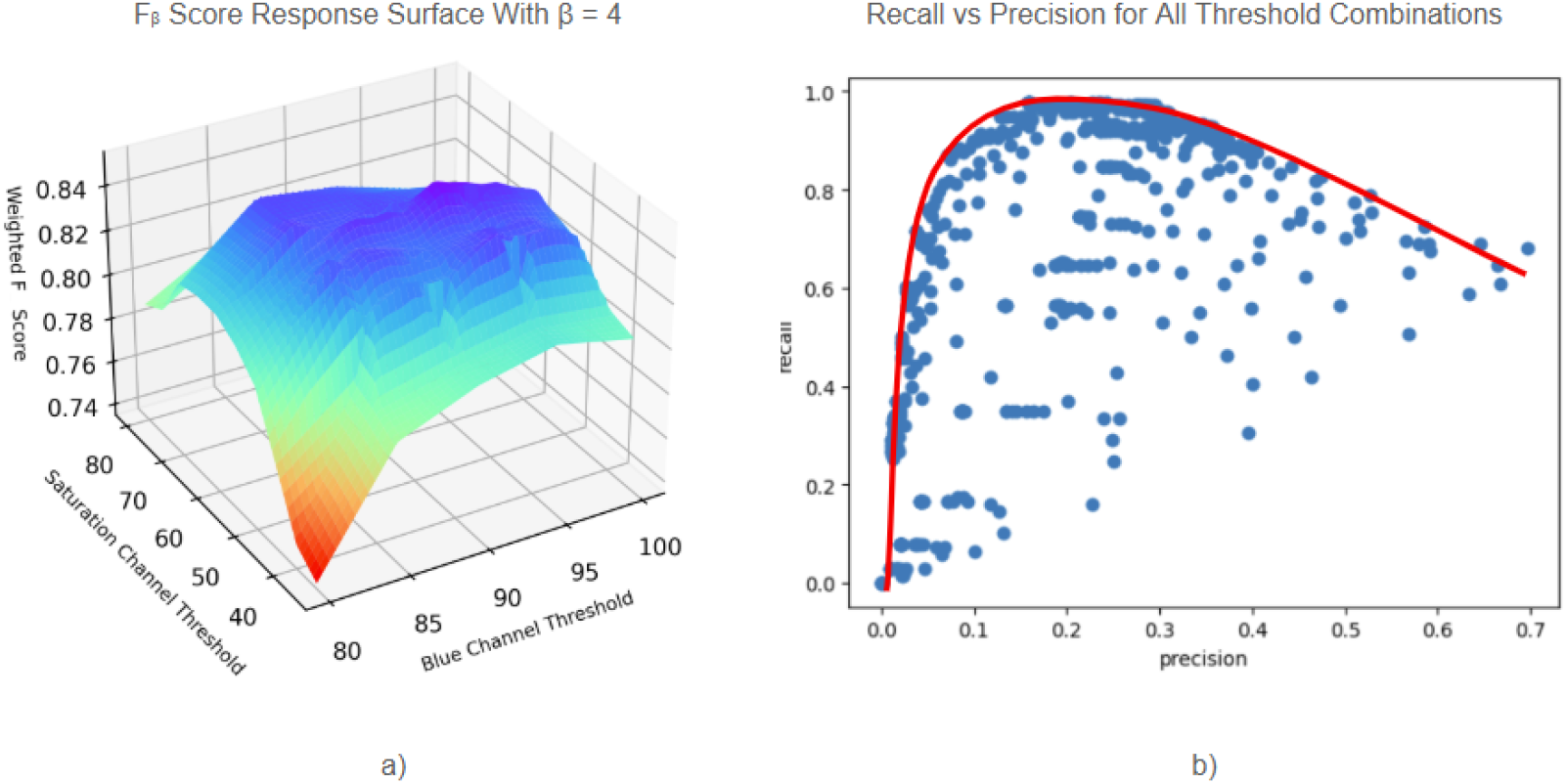
The F-score response surface (a) illustrates the balance between recall and precision for each model evaluated in the parameter grid search, highlighting the optimal threshold choice for a given β. The F-score combines precision and recall into a single value using β as a weighting factor, where β > 1 gives greater importance to recall. Precision, recall and F-score were computed for every combination of our two tuneable parameters, evaluated over our image subset. The subset consists of the 188 images containing seals supplemented with representative background images. The global maximum on the surface in (a) corresponds to the optimal threshold when β = 4. Similar surfaces can be generated for other β values. While β = 4 yields higher recall, it also results in a significantly higher human review burden. We do not treat β as an internal tuning parameter in this process, rather it is a final design choice dependent on both the desired application of the model and the available resources. (b) shows a scatter plot created by calculating the recall and precision for all threshold combinations. This provides an additional decision-making tool, aiding in the selection of the desired β based on constraints such as resource availability and required performance. Moving past the peak of the red frontier the gradient captures how quickly the recall falls off as you seek greater precision. Threshold choices beneath the frontier are strictly sub-optimal.

Table 2 presents the training and test recall for *β* =1, 2, 3, and 4 across the thinned dataset, where a higher *β* places increasing importance on recall. As a more insightful metric than precision in our semi-automated system with expert review, we present the search area reduction for the human reviewer across the dataset of >5,500 images (as precision was close to 100% for experts reviewing these high resolution images). The search area consists of the cumulative area covered by the clipped images of potential seals, as well as any complex images flagged for traditional manual review.

**Table 2.** Model performance metrics for varying values of β, showing recall on training and test datasets, and the percentage of image area requiring manual review. Higher β values favour recall over precision. A value of β = 2 was selected for the final model, balancing detection sensitivity with the human effort required for expert review over the >5,500 images in the 2023 WAP survey dataset.

| $\beta$ | Train Recall | Test Recall | Area Reduction |
| --- | --- | --- | --- |
| 4 | 97% | 99% | 90% |
| 3 | 93% | 96% | 92% |
| 2 | 83% | 82% | 94% |
| 1 | 68% | 68% | 95% |

The human review stage of this semi-automated pipeline was completed by an expert for the *β*=2 case, processing an equivalent of <6% of the total survey area. The combined recall of the automated and human review stages across the 800 manually reviewed images remained 82%.

### ii. Density estimation

The density estimates by species are summarised in Table 3. We used the presented algorithm to analyse a >5,500 image subset of the imagery captured in 2023 along the track lines in Figure 2. Using Equation 4, we computed a 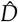 of 2.61 ice seals per km^2^. We identified a total of 758 unique seals. Of these 758 seals, manual species identification was carried out by an experienced observer with the species split between crabeater and Weddell’s being approximately even in cases where species could be confidently determined.

**Table 3.**
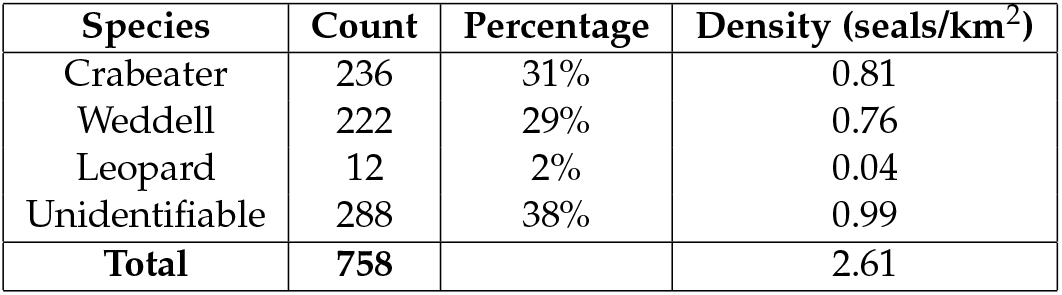
Total seal counts, species-level distribution, and estimated on-ice densities derived from the semi-automated analysis of 350 km^2^ of 2023 aerial imagery. Crabeater and Weddell seals made up the majority of identifiable individuals. Densities were estimated using a correction for model recall (Equation 4). Unidentified seals reflect cases where image resolution precluded confident classification.

| Species | Count | Percentage | Density (seals/km<sup>2</sup>) |
| --- | --- | --- | --- |
| Crabeater | 236 | 31% | 0.81 |
| Weddell | 222 | 29% | 0.76 |
| Leopard | 12 | 2% | 0.04 |
| Unidentifiable | 288 | 38% | 0.99 |
| **Total** | **758** |  | **2.61** |

## IV. Discussion

### i. Semi-automated pipeline

The semi-automated pipeline aims to reduce the human review time while improving recall by mitigating the inherent imbalance in the dataset via search area reduction. The unaided expert reviewer, who conducted traditional manual reviews over entire images, detected 63 seals in 800 images. The second expert, aided by a coarse filter, detected 63 additional seals. These 126 seals form a ground truth estimate, with an unknown number of seals potentially undetected by either method. The true recall for this unaided manual review is at best 50%, which forms the baseline against which we compare our new model performance. The manual review performance is reflective of the imbalance in the dataset [45] as well as the difficulty in identifying such small objects among a vast background (Figure 1); a skew similarly reflected in other studies [43, 20]. Our model’s improved recall of 82% stems in part from the re-framing of the question posed to the reviewer into an easier, more balanced task [45]. In traditional review, the reviewer must answer the question “How many seals are in this large landscape?”, a task which combines object localisation and object classification; in our method, the reviewer only classifies whether a bite-sized image is a seal or not.

It should be noted that the recall and precision of the model’s automated stage are distinct from the recall and precision of the overall method, which includes human review of the clipped images. The human can only discard clipped images. Thus, the overall method recall is less than or equal to the recall in the automated stage, with the difference depending on the human reviewer’s false negative rate. Likewise, the human quickly discards false positives served by the algorithm, increasing the method’s overall precision. Expert review of the high resolution clipped images had near perfect recall and precision. However, the decision to perform expert review in the *β* = 2 scenario reflects a bottleneck in expert availability to process the downstream clipped images. In future research, we intend to investigate the potential of both crowd sourcing this final review stage and completely replacing the human review with a second classification algorithm (*i*.*e*., [56]). We expect degradation of model performance when replacing expert review with either. However, outsourcing the final review stage to crowds or algorithms facilitates a higher *β* selection, which could potentially offer improved overall recall than our current 82%.

Our model systematically failed to detect seals in areas of dark shadow on the ice, likely due to both the lower contrast between the seal and its background and the elimination of large dark shadows that may contain these seals in the contouring stage. This was a rare occurrence in the labelled dataset with just three instances of seals in shadow. Methods to overcome this blind spot could include the incorporation of adaptive contrast enhancement [57] or other preprocessing techniques [58, 59]. However, with insufficient data points, there is higher risk that efforts to recapture these seals introduces overfitting. As more data is gathered, efforts to handle these edge cases can be more robustly developed.

Figure 7 displays a mosaic of seal proposals before the human review stage, with arrows added manually indicating true seal detections. Within these images, those not annotated with arrows, show examples of false positives, primarily stemming from geographical features in the ice. Investigating additional methods to remove these features could improve the precision of the automated stage and further reduce the necessary human input. One previous avenue that has been explored was the addition of infrared imagery to aid in the detection of penguins and pinnipeds in overhead drone imagery [60]. However, the addition of this data stream was not shown to increase model performance in their test case. Although our developed model will not generalize to other animals, environments, or conditions, the principle of taking a priors based approach to assist in processing large datasets is sound [61]. For each new problem, new priors must be defined and tuned, but the architecture can be recycled. Finally, it should be again noted that the ground truth in these datasets can only be estimated, as there is no way to guarantee seals are comprehensively detected. Any blind spots in the ground truth estimate may translate to blind spots in any model trained on the data.

**Figure 7.**
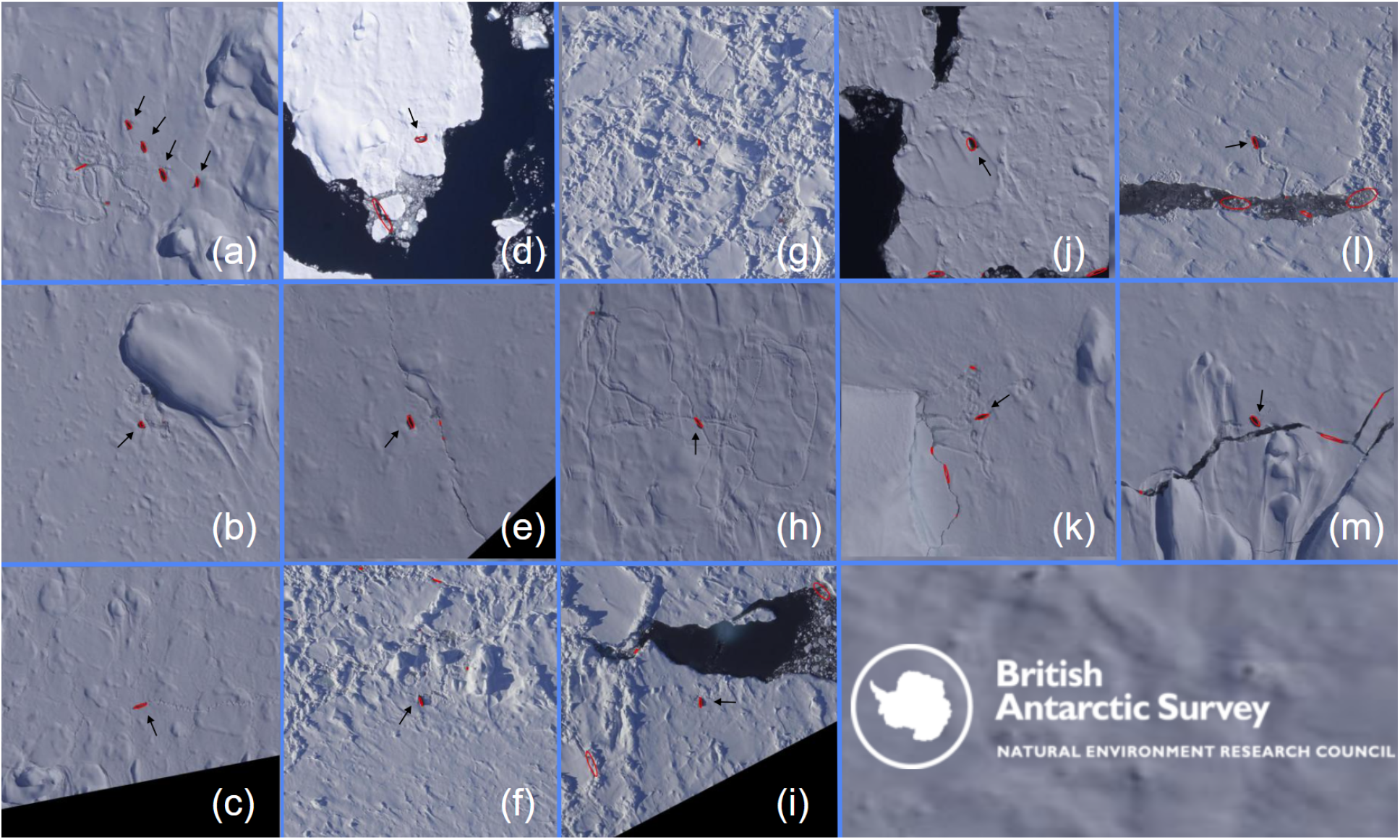
A collection of images from the 2023 survey showing typical model outputs. The areas circled in red have been identified by the model as potential seals. All true seal detections are indicated by a black arrow. All remaining model detections are examples of false positives, which must then be discarded by the human reviewer. False positives typically follow terrain features. Developing additional methods to remove terrain features could improve model precision.

### ii. Survey efforts in previous years

In 1999, observer-based aerial surveys were conducted as part of the UK’s contribution to the Antarctic Pack Ice Seals (APIS) surveys [62, 20]. These surveys were flown between the 22^nd^ and 29^th^ January 1999, during the moult period, where seals are hauled out on ice floes. These surveys specifically targeted areas of pack ice and, to date, provide the most comprehensive published assessment of the density and abundance of pack-ice seals on the Antarctic Peninsula. The stated crabeater seal density in the Marguerite Bay area was the highest of the areas observed at 8.428–20.258 seals/km^2^, with an abundance estimate of 373,132. The flight lines conducted during these surveys are shown in Figure 8 with red lines. However, the 1999 surveys differ in both scope and methodology from those conducted in 2023. Firstly, the 1999 survey covered a larger set of study areas, with data collected in real time by human observers with unknown recall and without comprehensive, permanent imagery that can be referred back to. Secondly, their estimates for seal density include corrections for seal haul-out proportions, a factor highly dependent on both environmental and behavioural conditions. In the absence of bio-logging data corresponding to our own survey dates in November, we cannot directly compare recorded densities.

**Figure 8.**
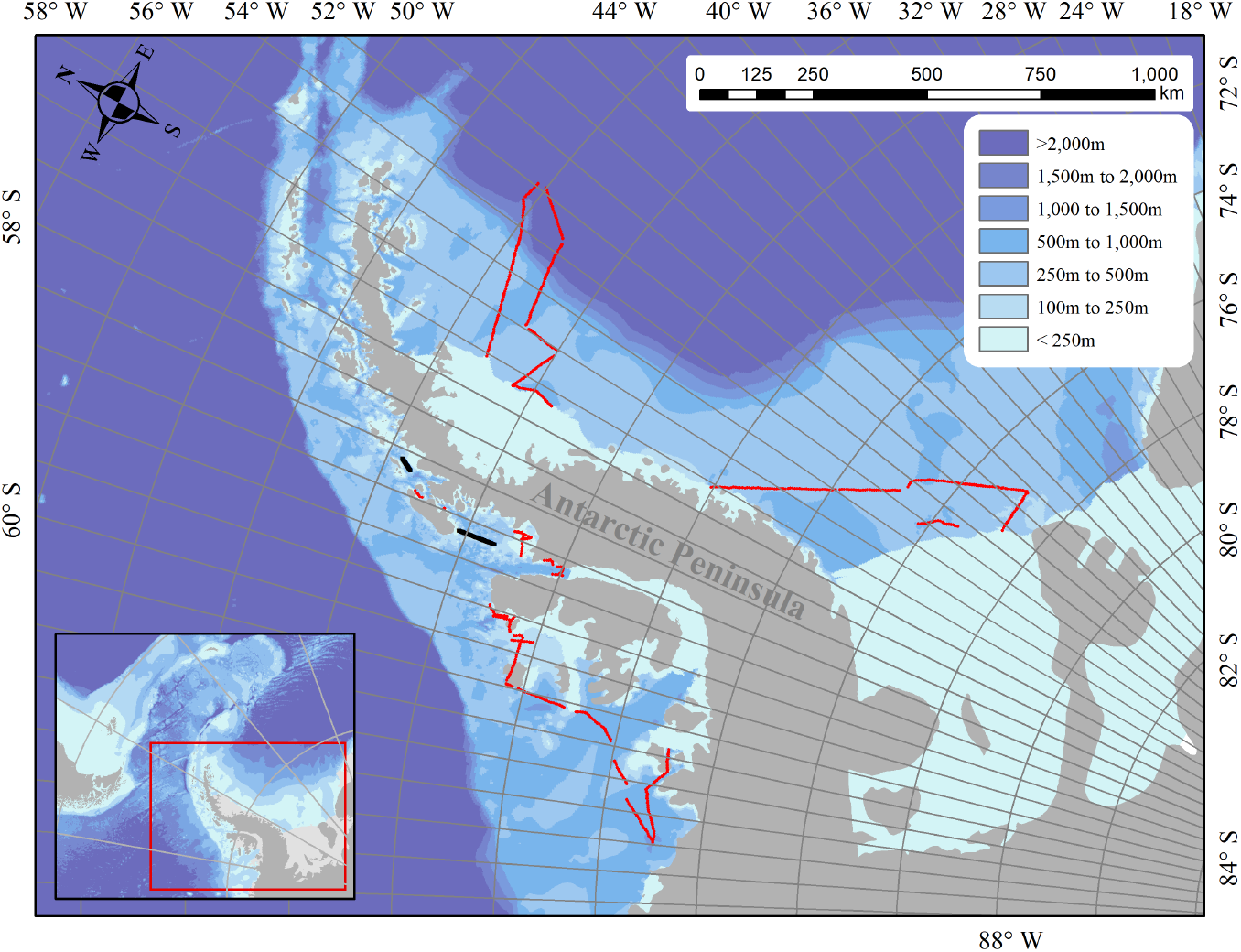
Previous ice seal aerial surveys conducted on the Antarctic Peninsula. Surveys conducted as part of the UK’s contribution to the Antarctic Pack Ice Seals (APIS) surveys in 1999 [62, 20] are depicted in red, with BAS aerial surveys flown in 2015 in black.

The next most significant aerial survey effort within the Crystal Sound and Marguerite Bay area consisted of 1041.6 km of flight lines in late October / early November 2015; Figure 8, black lines. These data are available on request from the UK Polar Data Centre. In the scope of this study, areas of sea ice (282.7km^2^) in the acquired imagery were manually counted ice seals by a single observer by loading the imagery into ArcMap v10.3 and placing shapefile points on top of every identified seal. Analysis of this imagery data produced a total of 800 seals, corresponding to a density of 2.83 seals per km^2^. Although no bio-logging data was utilised in this survey, we are again unable to compare our density estimates directly without introducing bias. Furthermore, as the imagery collected in 2015 was panchromatic, and therefore they lack the full spectral range of information needed to apply our semi-automatic pipeline.

Historically, the substantial time and effort required to manually process aerial imagery has limited the scope and regularity of aerial surveys targeting ice seals. As a result, survey efforts have been infrequent, methodologically inconsistent, or opportunistic, which has limited the availability of reference baselines. Table 4 outlines key differences in survey design, and Figure 8 shows the variation in spatial coverage across years. Together, these undermine the ability to detect true trends in seal populations. Overcoming these constraints is essential for generating robust, repeatable population estimates.

**Table 4.** Key methodological differences between the 1999, 2015, and 2023 seal surveys in the Western Antarctic Peninsula. Variations in timing, imaging platforms, tagging data, and image formats affect data comparability across years. Only the 2023 survey used a full-colour RGB camera compatible with the current semi-automated detection pipeline, underscoring the need for standardised future survey protocols. However, the lack of concurrent tagging data prevents extrapolation of on-ice densities into population estimates.

|  | 1999 | 2015 | 2023 |
| --- | --- | --- | --- |
| Survey Method | On-board observer | Panchromatic camera | RGB camera |
| Time of year | January | late Oct / early Nov | mid Nov |
| Concurrent tagging data | Y | N | N |
| Uncorrected counts available | N | Y | Y |
| Area calculation | unknown | ice only, water excluded | ice only, water excluded |
| Suitable for our pipeline | N | N | Y |

Our development of a semi-automated image analysis pipeline addresses this bottleneck, reducing the area requiring manual review by more than 95% and making it feasible to conduct standardised, high-resolution surveys at regular intervals. By enabling consistent data collection, this approach lays the groundwork for meaningful long-term monitoring of ice-obligate seal species in the WAP, or further afield. We strongly recommend the concurrent deployment of satellite-linked bio-logging instruments, where feasible, to complement surveys and enable abundance estimates rather than being limited to haul-out densities alone. Trans-dermal tagging campaigns, which mitigate against tag loss during the annual moult, are essential to quantify haul-out probabilities and correct for detection biases, providing the missing link needed to convert on-ice counts into robust population estimates.

The abnormally low sea ice conditions on the WAP in 2023 [48] may have contributed to unusually low observed densities of ice seals. While these trends could indicate future patterns if sea ice remains scarce, they should be interpreted cautiously until verified by further surveys and further sea ice monitoring. The unusually low sea ice across the WAP region may have prompted ice seals to redistribute away from historically abundant areas; yet, the presence of dense pack ice in the surveyed areas, which would typically attract seals, suggests the possibility of a more concerning decline.

## V. Conclusion

Monitoring ice seal populations is crucial, as through a more developed understanding of their population dynamics, we can gain a more developed picture of the wider health and functionality of the WAP ecosystem. This is particularly important in light of climate change and increased krill fishery pressure on the WAP [63]. However, the processing of increased volumes of primarily imagery-based survey data in line with the fast paced advances in remote sensing places onerous demands on researchers. Here, we present a semi-automatic model developed to process a >5,500 image subset of the imagery captured in 2023 during our aerial survey campaign over Marguerite Bay and Crystal Sound. By using a priors based approach and unsupervised machine learning, we sidestep several key barriers for deep learning: (i) insufficient volume of readily available labelled data; (ii) the imbalanced nature of the classification problem; (iii) and low quality ground truth estimates due to human error exacerbated by data volume and imbalance. We tuned our model using the F score with *β*=2 and achieved a recall of 82% after expert review of automatically generated image clippings, outperforming the ∼50% recall for unaided manual review, while reducing the survey area, and therefore human input, by >94%. In future research, we will evaluate both crowd sourcing and a secondary classifier to replace the expert review, enabling a higher *β* selection which could further improve the model’s overall recall.

While previous aerial surveys have offered valuable snapshots of seal presence along the WAP, methodological inconsistencies have limited their utility for assessing long-term trends. By providing a scalable, reproducible, and efficient framework for image-based survey processing, higher-frequency, standardised surveys become feasible. When paired with concurrent tagging campaigns, we can uncover population trends for key indicator species and take a critical step toward more responsive and evidence-based polar conservation policy.

## Author Contributions

Conceptualisation: CB, JF, PTF; Data collection: NF & AF; Analysis: CB & CM (lead on Machine-Learning approach); Manuscript drafting: CB & CM. All authors contributed to the final review of the manuscript.

## Code and Data Availability Statement

The code accompanying this research is openly available in github.

## Acknowledgments

The authors would like to acknowledge open source developers, especially Nima Moshtagh, whose Minimum Volume Enclosing Ellipsoid method from the MATLAB Central File Exchange was adapted for part of this work.

